# Robustness and lethality in multilayer biological molecular networks

**DOI:** 10.1101/818963

**Authors:** Xueming Liu, Enrico Maiorino, Arda Halu, Joseph Loscalzo, Jianxi Gao, Amitabh Sharma

## Abstract

Robustness is a prominent feature of most biological systems. In a cell, the structure of the interactions between genes, proteins, and metabolites has a crucial role in maintaining the cell’s functionality and viability in presence of external perturbations and noise. Despite advances in characterizing the robustness of biological systems, most of the current efforts have been focused on studying homogeneous molecular networks in isolation, such as protein-protein or gene regulatory networks, neglecting the interactions among different molecular substrates. Here we propose a comprehensive framework for understanding how the interactions between genes, proteins and metabolites contribute to the determinants of robustness in a heterogeneous biological network. We integrate heterogeneous sources of data to construct a multilayer interaction network composed of a gene regulatory layer, and protein-protein interaction layer and a metabolic layer. We design a simulated perturbation process to characterize the contribution of each gene to the overall system’s robustness, defined as its *influence* over the global network. We find that highly influential genes are enriched in essential and cancer genes, confirming the central role of these genes in critical cellular processes. Further, we determine that the metabolic layer is more vulnerable to perturbations involving genes associated to metabolic diseases. By comparing the robustness of the network to multiple randomized network models, we find that the real network is comparably or more robust than expected in the random realizations. Finally, we analytically derive the expected robustness of multilayer biological networks starting from the degree distributions within or between layers. These results provide new insights into the non-trivial dynamics occurring in the cell after a genetic perturbation is applied, confirming the importance of including the coupling between different layers of interaction in models of complex biological systems.

## I. INTRODUCTION

The recent development of high throughput omics technologies has facilitated the extensive profiling of the different molecular *strata* composing living organisms, such as the transcriptome, epigenome, and proteome, providing a more comprehensive picture of the detailed molecular composition of cellular systems. However, cellular processes are not only driven by individual molecules but also by the interplay between them. These interactions are conventionally modeled as context-specific molecular interaction networks [1], such as gene regulatory networks [2], protein-protein interaction (PPI) networks [3], and metabolic networks [4, 5]. Such network-based analysis [6] has become an effective and widely used tool in the analysis of cellular systems. While the study of the static topology of these networks has been successful in various applications, such as disease gene prioritization [7–9], disease biomarkers discovery [10], and disease diagnosis and subtyping [11], substantial insights can be gained by analyzing the properties of dynamical processes evolving over the nodes and edges of the network. These processes are usually defined to mimic the effects of environmental changes, internal perturbations, onset of diseases, or random failures occurring in the network [12].

An established way to quantify the effect of perturbations in biological systems is the analysis of their robustness, defined as their ability to maintain stable functioning despite various perturbations [13, 14]. In biological systems across all scales, from cells to organisms, robustness is attained by a combination of five mechanisms: feedback control, structural stability, redundancy, modularity, and adaptation [15, 16]. For example, by applying percolation theory to the analysis of the robustness of biological networks [17, 18], Jeong *et al.* [19] found strong connections between the centrality of a protein and its lethality; metabolic networks are exceptionally robust [20], hinting at why organisms can survive under different environmental conditions; robustness analysis of molecular networks under perturbations has become an efficient tool for uncovering disease mechanisms at the molecular level [21, 22]. Most of these studies focus on the investigation of single molecular networks. However, molecular networks are not independent, and processes can span multiple molecular layers simultaneously, generating intricate patterns that are difficult to uncover when networks are analyzed separately [12, 23]. For example, in a cell, genes can activate or inhibit other genes, and this regulation is operated through physical protein-protein and protein-DNA interaction bindings. Proteins can, in turn, affect metabolic reactions through physically or functionally associating with metabolites. In these cases, exploring networks of molecules of the same kind in isolation can ultimately lead to an incomplete or even incorrect picture of the problem. Thus, accounting for the interactions between different molecular networks is critical for understanding cell dynamics and functionality.

In network science, systems composed of multiple interacting networks [24–28] have attracted considerable attention owing to the discovery of novel structural and dynamical features that differ from those observed in isolated networks. In the past decade, the mathematical frameworks for characterizing the robustness of a network of networks [27] or multi-layer networks [28] have been studied in various settings, such as full interdependency [24], partial interdependency [29], interconnections [30], spatially embedded networks [31], multiple supports [32], directed networks [25], targeted attacks [33], multiple networks [34], and many more. The robustness of multilayer networks has a broad impact on infrastructure networks [35], ecological systems [36], social networks [37], and financial networks [38]. Recently, the growing availability of massive genomic, proteomic, and metabolomics data has stimulated the construction of multilayer biological molecular networks [39–44]. For example, Shinde and Jalan [45] proposed a multiplex network composed of six different PPI layers representing different life stages of *C. elegans*, showing varying degree-degree correlation and spectral properties across the nematode’s life cycle. Bennett *et al.* [46] found functional communities across layers in a two-layer PPI network of yeast, where one layer is connected by physical interactions and the other by genetic interactions. In this context, different layers model different kinds of interactions. Klosik *et al.* [47] designed a vast, directed, biological molecular network, called the interdependent network of gene regulation and metabolism, which is composed of three types of biological molecules: genes, proteins, and metabolites. For multicellular organisms such as humans, Didier *et al.* [48] and Valdeolivas *et al.* [49] investigated the community structure in multiplex biological molecular networks, which is composed of three or four biological networks sharing the same set of genes/proteins, with the nodes in each layer connected by different types of interactions, such as co-expression or physical interactions.

Despite advances in network theories and biological modeling as discussed above, our understanding of determinants of the robustness and lethality of multilayer biological molecular networks remains inadequate. The difficulty is rooted in four independent factors, each with its own layer of complexity:

1. *Biological modeling and data*. A comprehensive framework integrating heterogeneous sources of data of human molecular networks is still lacking. The integration of various molecular data, such as gene regulatory, PPI, and metabolic networks, is a challenging problem because these different components have completely different features and modes of interaction within an organism. Modeling these interactions in a meaningful way is critical for uncovering subtle molecular mechanisms responsible for disease pathogenesis and for identifying candidate genes for targeted therapies in precision medicine applications.
2. *Complexity in network topology*. The topological complexity of these interactions is another major challenge preventing us from understanding its functionality. Biological networks are of different kinds and the interdependence between them follow specific patterns. As an example, regulatory interactions between genes are usually modeled as directed, while PPI networks are undirected in virtue of their symmetric nature. How to meaningfully include such variety of interaction in a single description remains an open challenge.
3. *Diverse failure mechanisms*. We lack a reasonable failure mechanism to model how gene perturbations influence the function of the PPI and metabolic layers and the robustness of integrated multilayer biological networks. A perturbation of a gene may cause dysfunctions in its regulated genes and their protein products, while the dysfunction of proteins could cause other associated proteins to become non-functional since such proteins may be indispensable partners in the performance of cellular activities. Since proteins have functional associations with metabolites, the disruption of their functionality could affect the metabolic reactions they regulate. Although we cannot perfectly capture a holistic picture of the process by which a specific genetic perturbations propagates across a biological network, modeling the effect of gene perturbations with reasonable failure mechanisms could give us better understanding of the complex dynamics of the cell’s molecular machinery.
4. *Theoretical framework*. None of the previously described frameworks can be directly applied to multilayer biological molecular networks. Those approaches are, for the most part, agnostic of their applied setting and deal with networks of the same type, which are either all undirected [27] or all directed [25, 50], and in which where the interdependence relations are random. By contrast, biological networks include both directed and undirected network layers, and the non-randomly connected links between layers are of different types. In addition, the network sizes are different in scales. Thus, developing a general framework to analyze the robustness of multilayer biological networks remains an unsolved problem in interdependent networks [51].

Here, we discuss how to overcome these challenges and unveil the determinants of robustness and lethality of multilayer biological molecular networks, revealing the relationships between the critical structural features of a network and their functional importance in biology using a unified model.

## II. RESULTS

We start by integrating heterogeneous sources of data of human molecular networks, including a gene regulatory network, a PPI network, and a metabolic network. Further, we create a model for the robustness of a multilayer biological network, shown in section A. We validate the coupling between gene regulatory and PPI networks by comparing it to isolated networks in prioritizing the essential and cancer genes, shown in section B. In section C, we validate the relationship between lethality with metabolic disease genes. In section D, we compare the robustness of a multilayer biological network and its randomly rewired counterparts s, discovering that real networks are more robust. Finally, we developed a theoretical framework to analyze the robustness of multilayer biological networks.

### A. Model robustness of multilayer biological network

According to the central dogma of molecular biology, DNA is transcribed into RNA which is then translated into proteins. Many proteins can regulate metabolic reactions through physically or functionally associating with metabolites. Based on these well-known relationships, we constructed a multilayer network by aggregating the three major biochemical networks that govern cell function: a gene regulatory layer, a PPI layer, and a metabolic layer, as shown in Fig. 1.

**FIG. 1:**
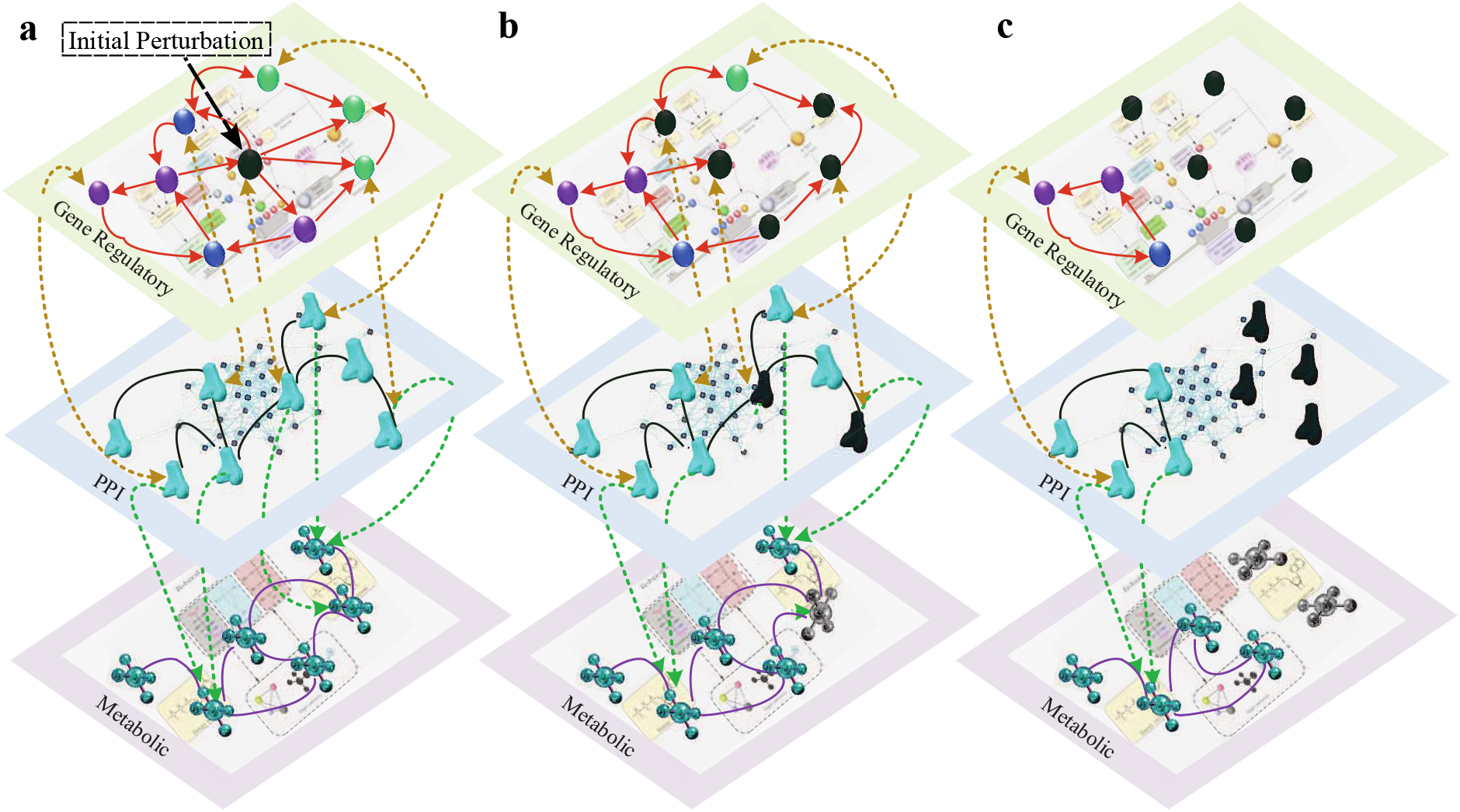
Schematic demonstration of the cascading failure process in the multilayer biological molecular networks. The multilayer model includes a gene regulatory network where the genes (ellipses) are linked by regulatory relations (orange directed links), a PPI network where proteins (bone shapes) are linked by physical interactions (green undirected links) and a metabolic network where metabolites (molecule shapes) are connected by chemical-chemical interactions (purple undirected links). The gene regulatory and PPI networks are connected by bidirectional interdependency links (yellow dashed lines). From the PPI to metabolic networks, there are multiple supporting links (green dashed lines). **a**. Initially perturb a gene in the gene regulatory network causing such gene stop functioning (represented by a black ellipse). **b**. The targeting genes of the perturbed genes fail (black ellipses), and their corresponding proteins stop functioning, represented by black bone shapes. **c**. The proteins that disconnected from the largest connected component fail (black bone shapes), and the metabolites losing its all supports from the PPI network stop functioning (black molecule shapes).

#### 1. Topology construction

We first construct three layers of biological molecular networks (the details for constructing the multilayer network are given in Methods section A):

1. *Gene regulatory network*. We use two types of gene regulatory networks in our work: a general gene regulatory network and three tissue-specific gene regulatory networks [22]. The general gene regulatory network is generated by curating the binding motifs of a subset of 695 human unique transcription factors, and the tissue-specific gene regulatory networks are curated from the FANTOM5 database [52].
2. *PPI network*. We combine several databases of physical protein-protein interaction data from high- and low-throughput experiments, obtaining a PPI network whose largest connected component consists of 15,906 proteins and 217,099 physical interactions.
3. *Metabolic network*. The metabolic network is constructed by curating the biochemical-biochemical (metabolite-metabolite) interactions from the STITCH database [53], and then mapping to metabolites in the Human Metabolome Database (HMDB) [54].
4. *Connections between Gene regulatory and PPI networks*. We connect the protein-coding genes in the gene regulatory network directly to their protein products in the PPI network. These connections result in 10,255 bidirectional interlayer links between the general gene regulatory network and the PPI network.
5. *Links from PPI to metabolites*. Protein-biochemical links are compiled from the STITCH database [55] and are directed from the protein to the metabolic layer as we make the simplifying assumption that the perturbation of an enzyme affects the metabolic reactions it regulates, while the opposite does not hold. Note that proteins and metabolites are connected in a many-to-many relation, since multiple enzymes and chemicals can participate in the same reaction, and multiple proteins can be associated with multiple metabolites. The interconnections between the PPI network and the metabolite network are obtained through the biochemical-protein links in the STITCH database. For the 15,906 proteins and the 1,269 metabolites in the multilayer network, we have 141,283 directed interlayer links connecting 12,039 proteins to 1,211 metabolites.

#### 2. Dynamical process on multilayer biological networks

To model the functionality and robustness of multilayer biological networks, we define a cascading failure mechanism simulating the effect of a perturbation in the network. From the molecular viewpoint, the cascade corresponds to a process whereby a number of perturbed transcription factors lose their ability to regulate their targets, resulting in some genes being left unregulated in the regulatory network, ultimately affecting the expression of the proteins for which they code in the PPI network. The altered expression of these proteins, in turn, disrupts the metabolic reactions they regulate.

The process is summarized below and shown in Fig. 1. Each node of these networks is assigned with a two-state variable, either “functional” or “dysfunctional”, and all the nodes are initially set as functional nodes. When a node becomes dysfunctional, it is removed from the network. The perturbation originates from a set of predefined target genes (TGs) in the gene regulatory network, simulating a loss of functionality due to e.g. mutations or gene knockouts. Since the TGs lose their ability to regulate other genes, both the TGs and the genes they regulate in the gene regulatory network become dysfunctional. As a consequence, the corresponding protein products of all of the involved genes become dysfunctional as well. After the removal of these proteins, the functional proteins that are left disconnected from the largest connected component of the PPI network become non-functional, along with their corresponding protein-coding genes. The regulated genes of the newly dysfunctional genes are then removed and the process continues in a cascading fashion until a stable state is reached.

In a metabolic network, each metabolite has multiple support links from the protein-protein interaction network. A metabolite stops functioning if a fraction *f*_P2M_ of its supporting proteins become dysfunctional, where *f*_P2M_ is a constant between 0 and 1. As an additional modeling choice, a metabolite is functional in any given time only if it belongs to the largest connected component. As shown in Fig. 1, the perturbation in the gene regulatory network can cause cascading failures propagating across the gene regulatory and PPI networks. When the process comes to a halt, the remaining nodes are identified as the final functional component.

### B. Perturbation on single genes revealing that the genes being important to system’s robustness enrich in essential and cancer genes

To investigate how the couplings between the gene regulatory and PPI networks contribute to defining the system structure and function, we compared the effects of the abovementioned perturbation process, hereby referred to as a “coupled” process, to the outcomes of a perturbation only affecting the PPI network alone, or an “uncoupled” process. The uncoupled process consists of removing the perturbed protein, their first neighbors, and any proteins that are left disconnected from the largest connected component thereafter. In order to provide a fair comparison between the two processes, we only consider perturbations of genes that are associated with their corresponding protein product, while genes that have no connections to the PPI layer are excluded from the analysis. In the PPI network of both uncoupled and coupled cases, we characterize the contribution of each node to the system’s integrity by measuring the final functional network size, 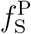, when that node is removed. A smaller final fraction of functional nodes indicates a larger contribution of the target node to system integrity. Thus, we assign an *influence* score 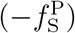 to each node to evaluate its influence on the system’s robustness, where 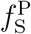 is the final functional size of the PPI network.

We compiled two sets of genes corresponding to biologically essential genes, from the Database of Essential Genes (DEG) [56], and cancer genes, from the Cancer Gene Census (CGC) [57]. Essential genes are indispensable for supporting cellular viability [56, 58], and cancer genes are those genes which contain mutations that have been causally implicated in carcinogenesis and that explain how dysfunction of these genes drives cancer development [57]. We calculated the precision-recall curves of coupled and uncoupled influence scores in recovering the sets of essential and cancer genes. As shown in Fig. 2a and b, the coupled influence scores yield higher precision-recall scores compared to the uncoupled and random scores, denoting the higher descriptive power provided by the layer couplings. In the coupled case, the removal of a single gene does not only cause one-time failures as those in the uncoupled cases, but also cause the second or even third round of cascading failures. we find that the averaged number of genes/proteins failed in the second round caused by the removal of a essential or cancer gene is higher than that of removing a non-essential or non-cancer gene, as shown in Fig. 2c and d, explaining why the coupled case performs better in prioritizing essential and cancer genes than the uncoupled case.

**FIG. 2:**
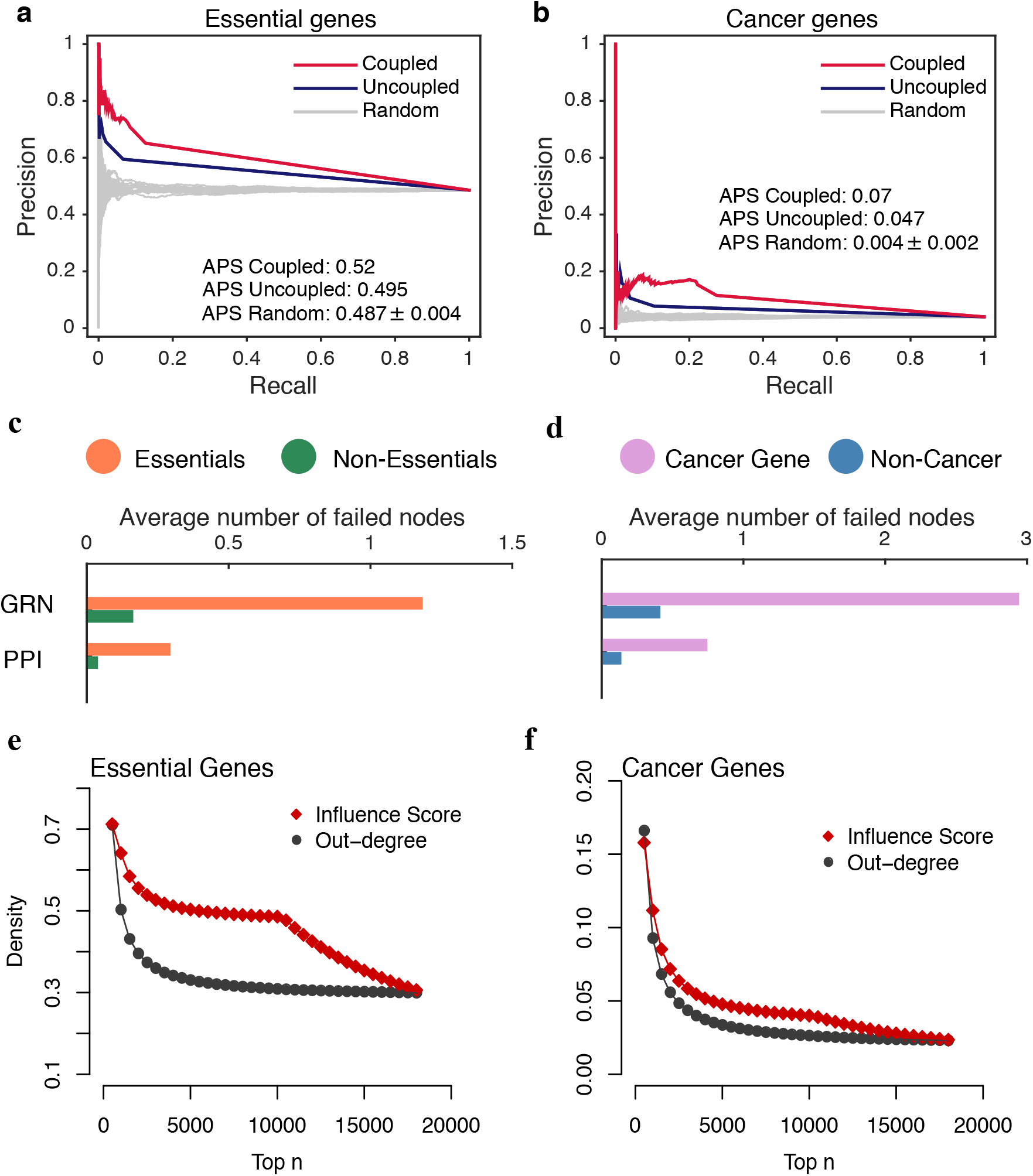
Comparison between the coupled and uncoupled cases. Average Precision-Recall (PR) curves of the coupled (red) and uncoupled (blue) influence scores in the prioritization of essential (**a**) and cancer (**b**) genes. The gray PR curves represent 100 random node rankings. On the right of each plot are listed the Average Precision Scores (APS) of the three ranking strategies evaluated from the corresponding PR curves. The performance of coupled influence scores in prioritizing essential and cancer genes are respectively 5.05% and 48.94% higher than that by uncoupled influence scores. In the coupled case, the removal of a single gene does not only cause one-time failures as those in the uncoupled cases, but also cause the second or third round of cascading failures. The average numbers of nodes failed in the second round caused by the removal of essential (**c**) and cancer genes (**d**) are respectively higher than that of removing the non-essential and non-cancer genes, explaining why the coupled case performs better in prioritizing essential and cancer genes. In addition, the densities of (**e**) essential and (**f**) cancer genes among the top n genes ranked by influence scores (red diamonds) are higher than that ranked by out-degrees (black circles). For the genes of the same influence scores or of the same out-degrees, we randomly put their orders 100 times, and compute the average densities. It indicates that the influence scores perform better than out-degrees in uncovering the connections between network topology and biological mechanisms.

Note that the same result holds when using different criteria for selecting essential genes: (1) probability of haploinsufficiency (Phi), (2) probability of loss-of-function intolerance (pLI), and (3) essential genes found by Dickinson et al. [59]. Genes with high scores of essentiality, as measured by these metrics, are associated to higher influence scores, and they are more prevalent among the genes with high influence scores in the coupled model compared with that of the uncoupled model (Figure S5). Note that the influence score in the coupled case incorporates the contribution of a gene to the integrity of the PPI network. We further test the performance of coupled influence scores in prioritizing disease genes categorized by their association with Mendelian or complex diseases. We divide the disease genes into MC (both Mendelian and Complex), MNC (Mendelian but Not Complex) and CNM (Complex but Not Mendelian) disease genes, as defined in [60]. The influence scores of genes show higher performance in prioritizing CNM disease genes (Figure S6), suggesting that complex disease genes have more cohesive connections to their surroundings. This observation aligns with the current understanding of complex diseases, which are hypothesized to stem from the interactions between a multitude of genes, requiring a higher degree of influence on their surroundings. By contrast, since Mendelian genes are, by definition, the primary cause of the disease phenotype they induce, their influence scores are indistinguishable from random chance.

Since the proposed failure mechanism depends on the out-degrees of the perturbed genes, we investigated to what extent the information provided by the influence score is different from the simple out-degree measure. We compute the fraction of essential, disease, and cancer genes among the sets of top *n* genes ranked by the influence scores and out-degrees, repeating the operation for each *n*. To account for the ambiguity in ranking caused by genes with the same out-degree or influence score, we randomly shuffled the ranks of the groups of genes corresponding to the same values 100 times, and computed the average ratios. As shown in Figs. 2 e and f, genes ranked by the influence score are enriched in larger fractions of essential and cancer genes compared to genes ranked by their out-degree, indicating that influence scores have a higher sensitivity in discerning the genes involved in critical cellular processes.

### C. Perturbation of a group of metabolic disease genes creates more damage to the metabolic network

Gene perturbations can propagate to the metabolic network through the failure of enzymes. For example, we consider the gene TCF7L2, one of the most replicated type 2 diabetes mellitus (T2D) susceptibility genes [61]. Its perturbation generates a cascade of failures that leads to the removal of o-hydroxyphenylacetic acid and lipid peroxidation in the metabolic layer. Lipid peroxidation has been observed to be directly associated with T2D [62], while o-hydroxyphenylacetic acid is formed from phenylalanine, an amino acid that is consistently associated with T2D risk [63]. As a more extensive experiment, we tested the outcomes of perturbing a group of metabolic disease genes in the gene regulatory network and observing the effects on the integrity of the metabolic network. We collected three sets of genes that are associated with dyslipidemia, blood pressure, and type 2 diabetes. From each set of metabolic disease genes, we randomly perturb a fraction *p* of genes and calculate the final functional metabolic network size, for *p* ∈ {0.1, 0.2, …, 1}. In order to control for node degree, for each metabolic disease gene set we generate a population of random gene sets of same size and similar degree as the original set, and repeat the perturbation process described above.

As shown in Fig. 3, and Figures S7-S12, perturbations targeting these gene sets cause more damage to the metabolic network with respect to degree-preserved random perturbations. For example, Fig. 3 shows the comparison between targeted perturbations (red boxes) of the dyslipidemia-related genes and random perturbations (blue boxes). In addition, we find that this result holds regardless of the value of the threshold *f*_P2M_ in the failure mechanism of the metabolic network (if more than a fraction *f*_P2M_ of supporting proteins fail, then the metabolite fails).

**FIG. 3:**
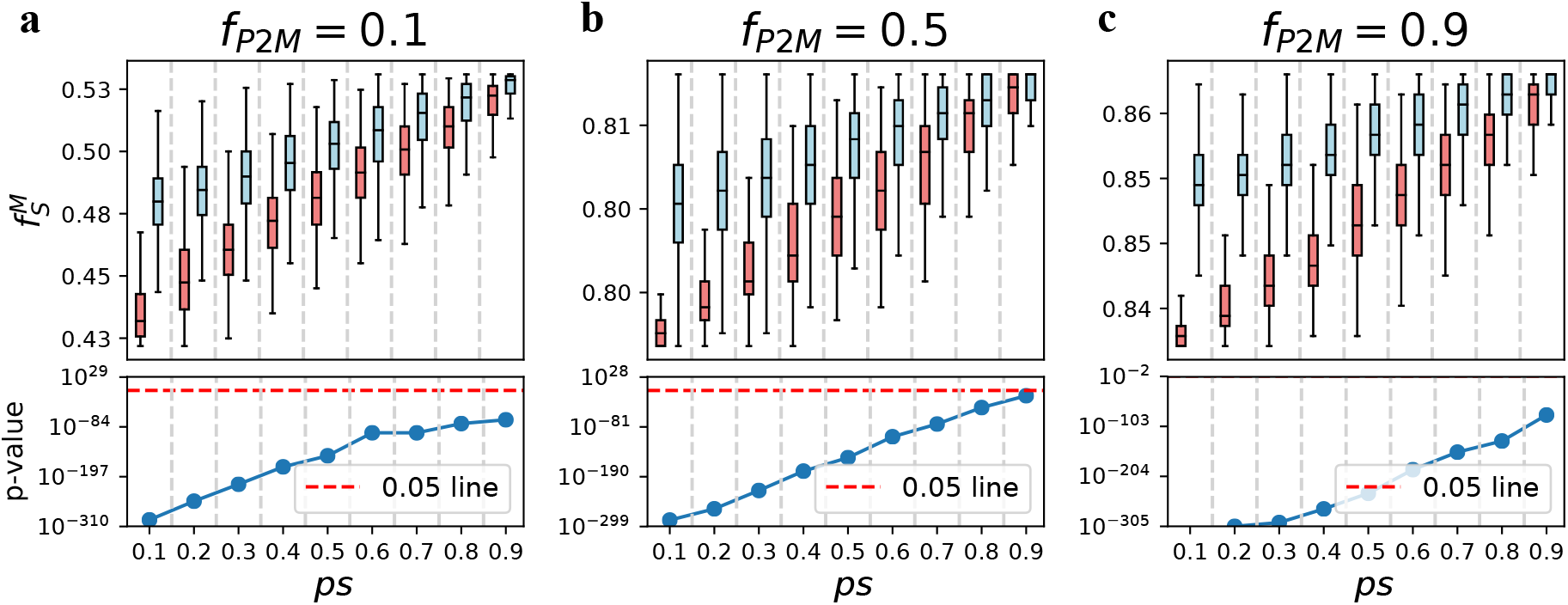
Targeting dyslipidemia-related genes (red boxes) causes more damages to the metabolic network than the degree-preserved random attacks (blue boxes). (top) Fraction of functional nodes after the perturbations for several values of remaining fraction *ps* of metabolic disease genes and threshold proportion *f*_P2M_ (a metabolite fails if more than *f*_P2M_ fraction of supporting proteins fail); (bottom) p-values of the Mann-Whitney test between the distributions of functional node set sizes in the targeted and random case. Lower values indicate a higher degree of damage to the network.

### D. Robustness of the multilayer biological network

For the real model of multilayer biological molecular networks, we could have multiple versions of randomized models: 1) intra-layer randomized versions, where the randomization can happen within the gene regulatory, PPI or metabolic layers. 2) inter-layer randomized versions, where the gene-protein or the protein-metabolite connections are randomly rewired. For each randomizations, it has three directions of rewiring methods: forcing the connections to be assortative or disassortative, or neutral without changing the degree correlations and degree distributions. Assortative randomization in a directed gene regulatory network is realized by keeping the in-degree and out-degree distributions unchanged but increasing the correlations between the in-degree and out-degree of each node, while that in the PPI or metabolic networks is achieved by rewiring the high-degree nodes to the high-degree nodes and low-degree nodes to the low-degree nodes, with their degree distributions unchanged. Assortative randomization in couplings involves connecting the high-degree nodes in one layer to the high-degree nodes in the other layer. Similarly, disassortative randomization is realized by decrease the in-degree and out-degree correlations or connecting the high-degree nodes to the low-degree nodes.

The robustness of the multilayer biological network can be valued by the final functional sizes in three layers (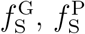, and we use 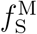 as a general metric representing any of them) after randomly removed 1 − *p* fraction of genes. The higher value of the final functional size indicates higher robustness. Or it can be valued by the integral size of functional network size 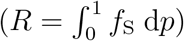, with *p* varing from 0 to 1. We find that the robustness of the multilayer molecular network is comparable to or higher than the robustness of the randomized models. For the disassortatively and neutrally randomized models, their robustness are comparable to the real models, as shown in Fig. 4**a** and Figures S13. We find that the real model is more robust than the following two randomized models: (1) randomization in the gene regulatory network keeping the in-degree and out-degree distributions and increasing the in-degree and out-degree correlations (Fig. 4 b and c); (2) randomization of the couplings between the gene regulatory and PPI networks keeping the degree distributions in gene regulatory and PPI networks unchanged but increase the degree correlations between the connected gene-protein pairs (Fig. 4 **d**).

**FIG. 4:**
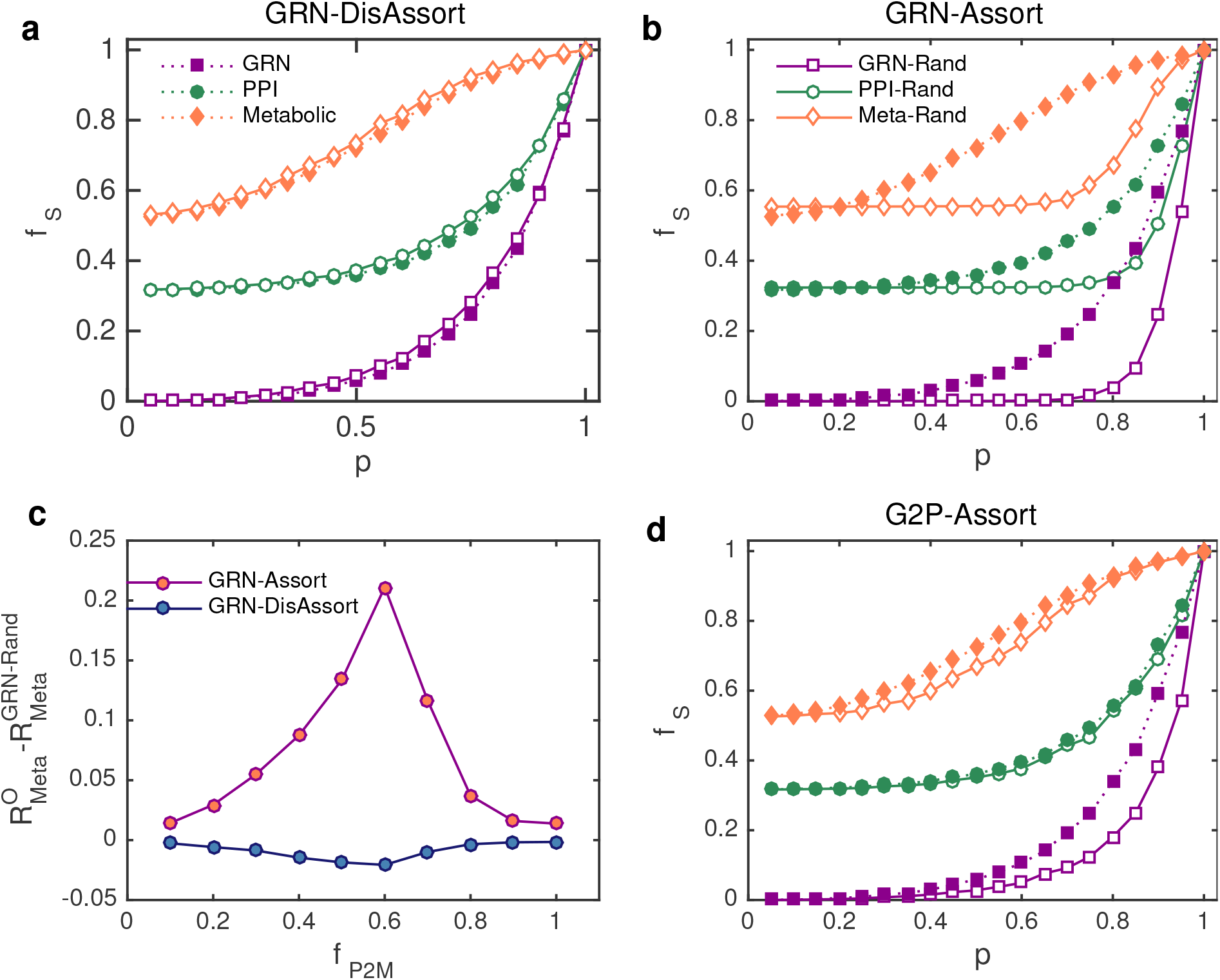
Comparing the robustness of the real biological multilayer networks (filled symbols) and randomized models (unfilled symbols). Higher values means higher robustness of the system. The real model has comparable robustness as **a.** the model that disassortatively randomized in the gene regulatory network, and higher robustness than **b.** the model that assortatively randomized in the gene regulatory network. Here the result in the metabolic layer is evaluated by setting the threshold *f*_P2M_ = 0.7, and the results under other threshold values are shown in **c**, where 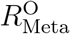 and 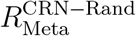 respectively represent the integral size of functional metabolic network in the real and randomized models, with *p* varing from 0 to 1. If the value 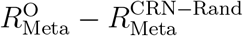 are larger than zero under different threshold, it means that the real model is more robust than the assortatively randomized model (GRN-Assort). The real metabolic layer has comparable robstness with the disassortatibely randomized model (GRN-DisAssort), since 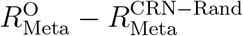 are near zero under different threshold. **d.** the real model is also more robust than the randomized model were the gene-protein connections are assortatively rewired.

### E. Comparison of numerical and analytical solutions

We first derive equations for calculating the functional network size in the gene regulatory network after initial perturbations on the 1 − *p* fraction of genes (see *Methods*). Then, the dysfunctional genes cause their corresponding proteins to stop functioning, and the functional network size in the PPI network can be calculated based on the generating function formalism [64] and percolation theory [24]. For most previous frameworks of robustness in multilayer networks, they are proposed under assumptions that the connections between layers are random [29] or follow specific patterns [39] in topology, which usually do not hold in real cases. In multilayer biological networks, the connections between the gene regulatory and the PPI networks are not purely random in topology, which makes it difficult to quantify the amount of failures that go from the gene regulatory network into the PPI network theoretically. We propose an analytical method to determine two equivalent coupling strengths, *q*_G_ and *q*_P_ to quantify the theoretical density of interconnections between omics layers (see Supplementary Information). By using these two coupling strengths, the connections between the gene regulatory and PPI networks can be treated as random in the theoretical calculation.

Next, we present the solution for the final functional network sizes step-by-step according to the cascading process between the gene regulatory network and the PPI network. At the final stage of the cascading process, the final fraction of functional nodes in the gene regulatory and PPI networks are, respectively, *ψ_m_* and *ϕ_m_* by using the framework as shown in Eq. 5. In the metabolic network, a metabolite node fails if a fraction of more than *f*_P2M_ of its supporting proteins fail. By applying percolation theory, the final functional node size of the metabolic network is 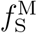 (see eq. 6 in *Methods*).

To verify the proposed framework, we first apply it to a synthetic model composed of three layers of Erdős-Rényi (ER) networks. We find that our framework accurately predicts the final functional network size in multilayer ER networks (see Supplementary file). Next, we repeat the calculations for the three-layer biological network described above. We evaluate the fractions of functional nodes at each stage of the perturbation, finding that the fractions of functional nodes in each cascading stage agree with the numerical simulations, as shown in Fig. 5**a**. To test the generality of our framework on multilayer biological networks with arbitrary degree distributions, we sequentially replace the gene regulatory layer with three tissue-specific gene regulatory networks, namely forebrain (Fig. 5**b**), lymphocytes (Fig. 5**c**), and lung (Fig. 5**d**). We find that in all three cases, our analytical framework correctly predicts the final sizes of functional nodes in these multilayer molecular networks.

**FIG. 5:**
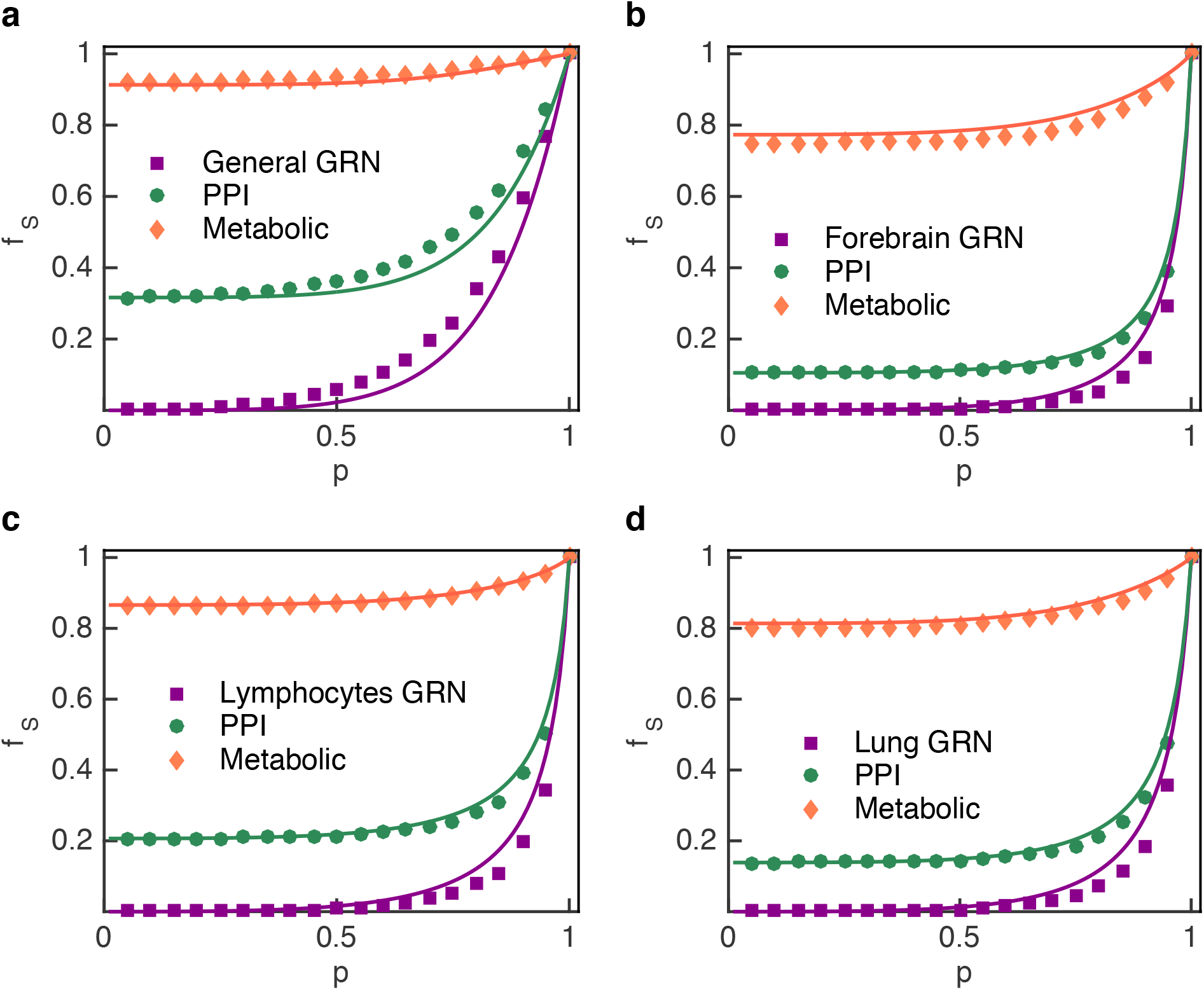
The final functional node sizes in the multilayer molecular networks after randomly removing 1 − *p* fraction of genes from the gene regulatory network. The gene regulatory network (GRN) in a is the general one, and those in **b, c** and **d** are the tissue-specific ones. The PPI and metabolic networks in these four panels are the same and a metabolite fail when all of its supports fail, that is *f*_P2M_ = 1. The theoretical results (solid lines) match the simulation results (symbols).

## III. DISCUSSION

To uncover how the couplings between molecular networks influence their biological functions, we propose a minimalistic model of multilayer molecular networks encompassing regulatory, protein-protein, and metabolic interactions, and develop a theoretical framework for analyzing the system’s robustness. We define a perturbation process that roughly simulates the cascade of effects occurring in the network when a group of genes is perturbed. We show that our analytical formulation correctly predicts the size of functional nodes in each stage of the cascading process. In this framework, we find that the topology of the proposed multilayer network is more robust than that of the randomized models. This finding could suggest that molecular networks may have evolved to avoid developing strong degree-degree correlations, as to increase the system’s robustness under perturbation.

We define an influence score characterizing the contribution of each gene to the system’s robustness, and find that essential and cancer genes are enriched in higher scores compared to random chance. In addition, to assess the contribution of the connections between different molecular layers, we compare the results above with the effects obtained by a perturbation process acting only on the isolated PPI network, finding that the multilayer system achieves superior performance in prioritizing essential and disease genes. Furthermore, targeting a group of metabolic disease genes causes significantly more damage in the metabolic network when compared to random perturbations, denoting the non-trivial association between these genes and the metabolic processes they regulate. The results above are complementary. On the one hand, we find that the coupled perturbation process accurately predicts genes that are important for biological processes and survival; on the other hand, we find that perturbing biologically important genes, defined *a priori*, causes more damage to the overall system’s integrity than perturbing other randomly chosen genes. The results in this work offer a comprehensive framework integrating heterogeneous sources of data of human molecular networks, opening new avenues to deepen our understanding of human molecular systems and their dynamics. Future directions of this work are two-fold: at a theoretical level, an important unmodeled factor in the analytical formulation is the correlation between in- and out-degree of the gene regulatory layer; at the biological level, there are additional molecular mechanisms that have major roles in determining the robustness of a cellular system, such as gene methylation and non-coding RNA regulation.

One additional avenue of investigation for improving this model is the inclusion of detailed molecular interaction parameters within the network, such as protein binding affinities and reaction rate constants, allowing for a generalization of the methodology to weighted networks. However, this approach has the drawback on relying on noisy and vastly incomplete sources of data, due to the lack of available information on most known interactions. Furthermore, the inherent study bias in these kind of measurements has the potential of generating spurious patterns that may confound the analysis. However, as high-throughput techniques for molecular profiling are developed and become more feasible, modeling these aspects can provide a deeper understanding of how perturbations spread in a heterogeneous biological interaction network.

## IV. METHODS

### A. Reconstruction of three layers of biological molecular networks

#### 1. Reconstruction of gene regulatory network

For the general regulatory network, a subset of 695 human transcription factor motifs, corresponding to 695 unique transcription factors, was curated from the list provided by an online library of transcription factors and their binding motifs [65]. For each of these 695 motifs, the entire hg19 genome was scanned using a program that scans sequence databases to find occurrences of known motifs [66], and significant hits with *p* < 1*e* − 3 were retained. 694 of the motifs had at least one significant hit in the genome for this scan. Once the genome-wide scan was completed, we took hg19 RefSeq annotated transcription start sites (TSS) and selected all associated Gene Symbols that mapped to a unique TSS. We then took the locations of the motif hits from the FIMO scan described above, and found the distance from the middle of the motif to the nearest TSS. Finally, we queried each of these files to find only motif-hits that occur in the promoter, where we defined the promoter as [−1000, +500] around the TSS. We used a p-value cutoff of 1e-6 for the regulatory network layer of our multilayer network, which results in 18,566 nodes and 65,310 links.

The tissue-specific gene regulatory networks are reconstructed by integrating transcription factor (TF) sequence motifs with promoter and enhancer activity data from the FANTOM5 project [52]. We curate tissue-specific regulatory networks using the FANTOM5 database [52], using the smaller “Network compendium” dataset, and select three tissue-specific regulatory networks, “forebrain” (ID:03), “lymphocytes” (ID: 12), and “lung” (ID: 23). These regulatory networks are downloaded from the website http://regulatorycircuits.org, and we set an link weight threshold of 0.05.

#### 2. Construction of protein-protein interaction network

For the protein-protein interaction network, we use physical protein interactions with experimental support, and do not include interactions extracted from gene expression data or evolutionary considerations. In order to obtain an interactome as complete as currently feasible, we combine several databases with various types of physical interactions:

- Regulatory interactions obtained from the TRANSFAC database [67]. Here nodes represent transcription factors, and connections represent physical binding to regulatory elements.
- Binary interactions: We combine several yeast-two-hybrid high-throughput datasets with binary interactions from IntAct [68] and MINT [69] databases.
- Manually-curated interactions from literature. We use IntAct, MINT, BioGRID [70], and HPRD [71] databases.
- Protein complexes: Protein complexes are single molecular units integrating multiple gene products. We use the CORUM [72] database, which is a collection of mammalian complexes derived from a variety of experimental tools, from co-immunoprecipitation to co-sedimentation and ion exchange chromatography.
- Metabolic enzyme-coupled interactions: Two enzymes are assumed to be coupled if they share adjacent reactions in the KEGG [73] and BIGG [74] databases.
- Kinase-substrate pairs: Protein kinases are important regulators in different biological processes, such as signal transduction using PhosphositePlus [75].

The union of all interactions yields a network of 15,966 proteins that are interconnected by 217,150 physical interactions. In this work, we focus on the largest connected component of the PPI network, which consists of 15,906 proteins connected by 217,099 links.

#### 3. Construction of metabolic network

For the metabolic network we use the STITCH database [53], which is an extensive association database that has both biochemical-biochemical (metabolite-metabolite) and biochemical-protein links. PubChem ids are used for metabolite identification, which maps well to metabolites in the Human Metabolome Database (HMDB) [54], facilitating the identification of metabolites. We limit our use of the dataset to interactions with experimental, similarity, and database evidence. The resulting metabolic association network, which we construct by combining the STITCH and HMDB databases, contains 1,398 metabolites with HMDB ids and 16,032 interactions between them, and its largest connected component include 1269 metabolites and 16019 links.

### B. Develop the theoretic framework for analyzing the robustness of multilayer molecular networks

We develop a general theoretic framework for modelling the cascading failures between the gene regulatory and PPI networks, and computing the final functional node sizes in three molecular layers after randomly removing a 1 − *p* fraction of genes from the gene regulatory network. We find that the gene regulatory network is vulnerable to perturbations, while the robustness in metabolic network is highly dependent upon the supports from the PPI network.

#### 1. Percolation analysis in single gene regulatory networks

We denote the joint degree distribution of the top-layer gene regulatory network with *P*_Gene_(*k*_in_, *k*_out_). We randomly choose a fraction 1 − *p* of nodes as “perturbed” genes. The probability density of genes with in-degree *k*_in_ and out-degree *k*_out_ not being perturbed or a targeting gene of one perturbed gene is *P*_Gene_(*k*_in_, *k*_out_)*p*^*k*_in_+1^. Thus, after removing the perturbed genes and their targets, the fraction of remaining nodes is

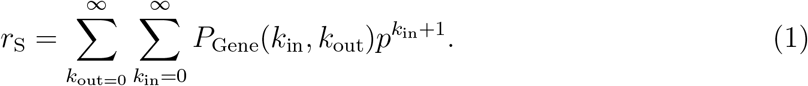

Assuming that there are no correlations between the in-degrees and out-degrees, the degree distribution of the remaining network can be written as

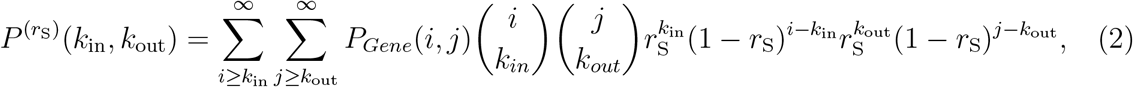

where 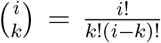, is a combination. In the remaining network, the isolates fail and the functional network size (described in detail in Methods) is

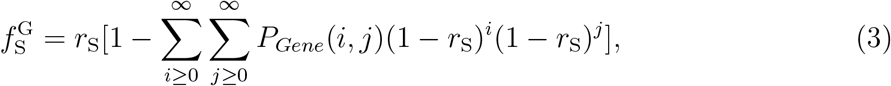

which can be simplified as

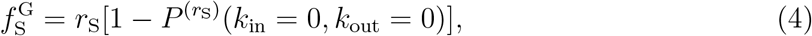

We apply this theoretical tool to gene regulatory networks including a generic network and three tissue-specific networks. As shown in the Figure S2 in the Supplementary Information, the theoretical results (solid lines) agree well with the simulations (symbols), confirming our theoretical analysis.

#### 2. Percolation analysis in coupled gene regulatory, PPI and metabolic networks

Owing to the incompleteness of the data, some proteins do not have corresponding genes in the regulatory networks, and some genes do not have corresponding proteins in the PPI network. Thus, the gene regulatory and the PPI networks are partially interdependent, and their interdependency relations are not random. We propose a method to find the equivalent coupling strengths between the gene regulatory and PPI networks, denoted by *q*_G_ and *q*_P_, so that the non-randomly interdependent relations could be approximated by random interdependency (see Supplementary Information).

We present the solution of the final functional network sizes step-by-step according to the cascading process between the gene regulatory and the PPI networks. In order to unify the quantities at each stage t of the cascading process, we define 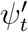 as the remaining network size after initial perturbation or receiving the failure from the PPI network, and *ψ_t_* as the functional network size in the gene regulatory network. At the initial stage *t* = 1, the remaining network size after perturbation is 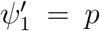, and the functional network is 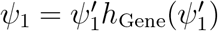, where *h*_Gene_(*p*) = *r_S_*/*p*[1 − *P*^(*r*_S_)^(*k*_in_ = 0, *k*_out_ = 0)]. Since a fraction of qp nodes in the PPI network depends on nodes from the gene regulatory network, the number of nodes in the PPI network become dysfunctional is 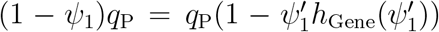. Accordingly, the remaining network size in the PPI network is 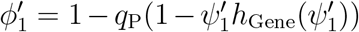. In the PPI network, the generating functions of the degree distribution and branching process are, respectively, 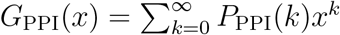 and 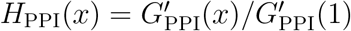. The fraction of nodes belonging to the largest connected component in the PPI is 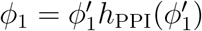, where *h*_PPI_(*p*) = 1 − *G*_PPI_(*px_c_* + 1 − *p*) with *x_c_* = *H*_PPI_(*px_c_* + 1 − *p*). Following this approach, we can construct the sequence for the remaining network sizes 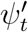 and 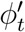, the functional network sizes *ψ_t_* and *ϕ_t_*. The general form is given by

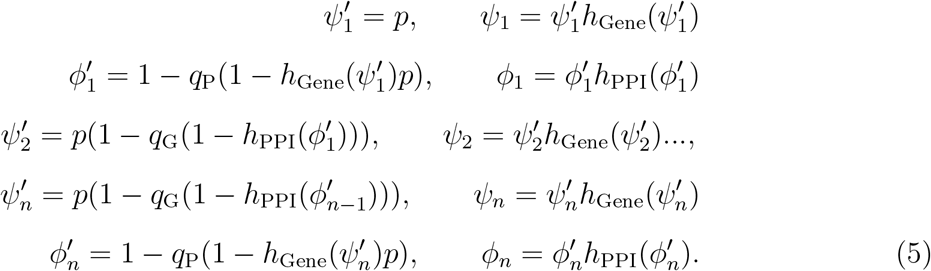

At the end of the cascading process, no further failures occur. The remaining fractions of nodes in the gene regulatory network and the PPI reach stable values 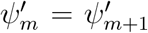 and 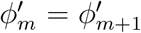 respectively. Thus, the fractions of the final functional nodes in the regulatory and PPI networks are, respectively, 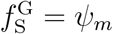 and 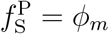.

The PPI and the metabolic networks are connected by unidirectional multiple support-dependence relations. In the metabolic network, *q*_Meta_ fraction of metabolites have multiple supports from the PPI network. The generating functions of the degree distribution of the metabolic network and its branching process are 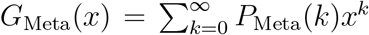 and 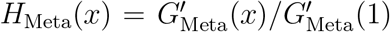, respectively. Each metabolite has *k_s_* supporting links from the PPI network, and we define the support degree distribution as *P*_D_(*k_s_*) whose generating function is 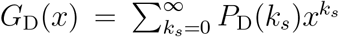, and whose branching process is 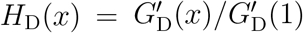.

Since the failure in the metabolic network cannot affect the gene regulatory and PPI networks, their percolation behaviors are equivalent to that in the coupled gene regulatory and PPI networks, whose final functional node sizes are *ψ_m_* and *ϕ_m_*, respectively. In the metabolic network, a metabolite node fails if more than f_P2M_ fraction of its supporting proteins fail. The probability that more than *f*_P2M_ fraction of the supports to a metabolite fail is 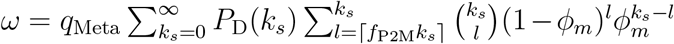. Thus, the fraction of the remaining nodes is *r*_Meta_ = 1 − *ω*. Thus, the final functional nodes size of the metabolic network is

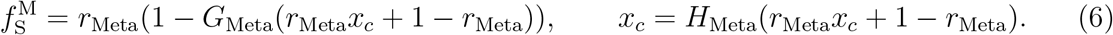

